# Maternal and cross-stage effects of *Metarhizium* fungal infection on the malaria mosquito *Anopheles coluzzii* life-history

**DOI:** 10.1101/2025.05.14.654141

**Authors:** Issiaka Sare, Mafalda Viana, Abel Millogo, Doubé Lucien Lamy, Assita Tenetao, Athanase Badolo, Florencia Djigma, Abdoulaye Diabate, Francesco Baldini, Etienne Bilgo

## Abstract

**Background:** Entomopathogenic fungi of the *Metarhizium* genus are widely used as biocontrol agents against harmful insects. These fungi are cost-effective and eco-friendly for vector control, providing an alternative to synthetic chemical insecticides. They have great potential as larvicides against malaria vectors, but their impacts on mosquito fitness have not been fully measured. This study evaluated the effect of *Metarhizium* fungal strains, locally isolated in Burkina Faso, on the larval survival of the mosquito *Anopheles coluzzii*, the life history of the emerging adults and on the maternal effects of exposed females.

**Methods:** We assessed the efficacy of *Metarhizium pingshaense* (strains S10 and S26) conidia against *An. coluzzii* larvae. First, larvae were reared in the presence of fungal spores and the survival of the larvae and adults that emerged, their wing length, oviposition rate and blood feeding behaviour were measured. Additionally, we assessed the efficacy of fungal strains S10 and S26 conidia against *An. coluzzii* adults by spraying them with spore suspensions and assessing the survival of their larval offspring. Survival data was analyzed using Cox proportional hazards model, while other life history traits using generalized linear mixed models.

**Results:** The fungal suspension applied to the water in which the larvae were reared caused mortality at the pupae stage. Only a small number of larvae emerged to reach adulthood. Furthermore, at the adult stage, these mosquitoes exhibited reduced survival compared to the control. However, body size and blood-feeding behavior were not affected by the treatment. When fungi was applied to adult females, the number of eggs layed was more abundant in infected group compared to controls, however a lower proportion of larvae successfully developed into adults.

**Conclusion:** The results of this study demonstrate the potential of *Metarhizium pingshaense* conidia for mosquito larval control. The identified cross-stage and maternal effects showed additional virulent effects of *Metarhizium*, thus reinforcing the evidence that this biocontrol agent should be part of an integrated vector management. Future work should focus on the molecular mechanism of the fungal infection at the larval stage to improve formulation or genetically engineer the conidia of these strains to make them more virulent.

## Introduction

The widespread use of insecticide-treated nets at the turn of the century was associated with a marked reduction in malaria mortality[1]. However, in recent years, the decline in malaria cases has stalled, threatening current malaria control efforts[2]. The increasing spread of insecticide resistance in malaria vector populations is among the factors contributing to the slowdown of the control efforts [3,4]. Indeed, the vast majority of vector control tools still rely on insecticide-based interventions, such as long-lasting insecticidal nets (LLINs) and Indoor Residual Spraying (IRS) so new and complementary tools will be needed for effective control. One tool gaining traction against malaria vectors are larvicides, or more broadly, tools that target the larval stage of mosquitoes [5]. Since larvicides are typically applied to larval breeding sites in the environment, traditional chemicals, while widely used, pose significant ecological risks due to their toxicity to non-target organisms and potential environmental persistence[6]. This has led to an increasing demand for alternative biocontrol solutions that are both selective and biodegradable. Among these, entomopathogenic fungi have emerged as a promising class of bio-insecticides with a narrower spectrum of action and reduced environmental impact[7]. These fungi, such as *Metarhizium anisopliae* and *Beauveria bassiana*, have demonstrated their ability to be specific to infect and kill larvae of major mosquito genera, including *Anopheles, Aedes*, and *Culex* [8]. Their mode of action involves the production of virulence factors and active metabolites that facilitate host invasion and ultimately lead to mortality. These metabolites also play a role in insect defense mechanisms against pathogens, further influencing host-pathogen interactions[9]. Experimental studies have provided substantial evidence supporting the efficacy of entomopathogenic fungi (EPF) in adult mosquito control, highlighting their potential as biopesticides in integrated vector management[10]. *Metarhizium* could also be effective for larval mosquito control because its spores can persist in aquatic environments, potentially infecting larvae through contact with contaminated surfaces or by ingestion, disrupting their development and increasing mortality.

Many laboratory and semi-field investigations have demonstrated that entomopathogenic fungal species such as *Metarhizium anisopliae* (ICIPE-30) and *Beauveria bassiana* (IMI-391510) exhibit significant larvicidal activity against major malaria vectors, including *Anopheles stephensi* and *Anopheles gambiae* [11]. These fungi act as natural biological control agents, infecting mosquito larvae primarily through direct contact with conidiospores present in the aquatic environment. Upon contact, the fungal spores adhere to the larval cuticle and germinate, forming specialized structures called appressoria, which facilitate penetration of the cuticle[12]. Once inside the host, the fungus proliferates within the hemocoel, disrupting physiological processes and leading to systemic infection[13,14]. As the fungal hyphae spread, they deplete larval energy reserves, produce toxic secondary metabolites, and compromise immune defenses, ultimately causing mortality[15]. The speed and efficacy of fungal infection depend on environmental factors such as temperature, humidity, and the larval developmental stage[12,16]. Additionally, some studies suggest that fungal infections may weaken larvae, making them more susceptible to other stressors, such as predation and chemical insecticides[17,18]. These findings highlight the potential of *Metarhizium anisopliae* as promising candidates for integrated vector management strategies targeting malaria-transmitting mosquitoes. Further studies have revealed that EPF target critical physiological systems within mosquito larvae. Histopathological analyses indicate that fungal infection disrupts the integrity of the cuticle, allowing fungal hyphae to invade internal tissues such as the alimentary and respiratory tracts. This invasion not only impairs nutrient absorption and metabolic functions but also leads to systemic mycosis, ultimately resulting in larval death[11,19]. Additionally, certain isolates of *B. bassiana* have demonstrated efficacy against culicine mosquito larvae (*Culex* spp.), suggesting a broader spectrum of activity across different vector species [20].

Beyond their direct lethal effects, EPF exhibit sublethal impacts that may further contribute to mosquito population suppression. Infected larvae often experience developmental delays, reduced pupation rates, and compromised adult emergence, all of which can disrupt population dynamics and transmission potential[21,22]. These multifaceted mechanisms position EPF as a promising alternative to conventional larvicides, particularly in the context of insecticide resistance and environmental sustainability. By offering an eco-friendly, target-specific, and potentially self-propagating solution, entomopathogenic fungi represent a valuable component of integrated vector management strategies aimed at reducing malaria transmission in endemic regions. These different studies conducted in the laboratory give satisfactory results on the ability of entomopathogens to control mosquito larvae. Our previous studies have shown that *Metarhizium* strains S10 and S26 two strains of fungi isolated in Burkina Faso in west Africa can reduce the survival of adult mosquitoes [23,24]. However, the ability of these two strains of *Metarhizium* to control mosquito larvae and the implications for the general fitness of the mosquito derived from the fungal suspension remain unknown. The aims of the present study were to investigate the potential use of the fungi *Metarhizium pingshanse* strains S10 and S26 to control both effect on larvae and subsequent emerging adults. This study evaluated the effects of larval exposure to entomopathogenic fungi on mosquito survival, development time, and adult emergence. We further assessed key adult life history traits, including feeding propensity, fecundity, and wing size as a proxy for fitness. The aim was to determine the cumulative impact of fungal exposure on mosquito biology and potential vectorial capacity.

## Methodology

We conducted two experiments. In the first we tested the cross-stage effect of infecting larvae with *Metarhizium* (**Figure 1A**). In the second, we exposed adults to *Metarhizium* and monitors the progeny life history traits.

**Figure 1:**
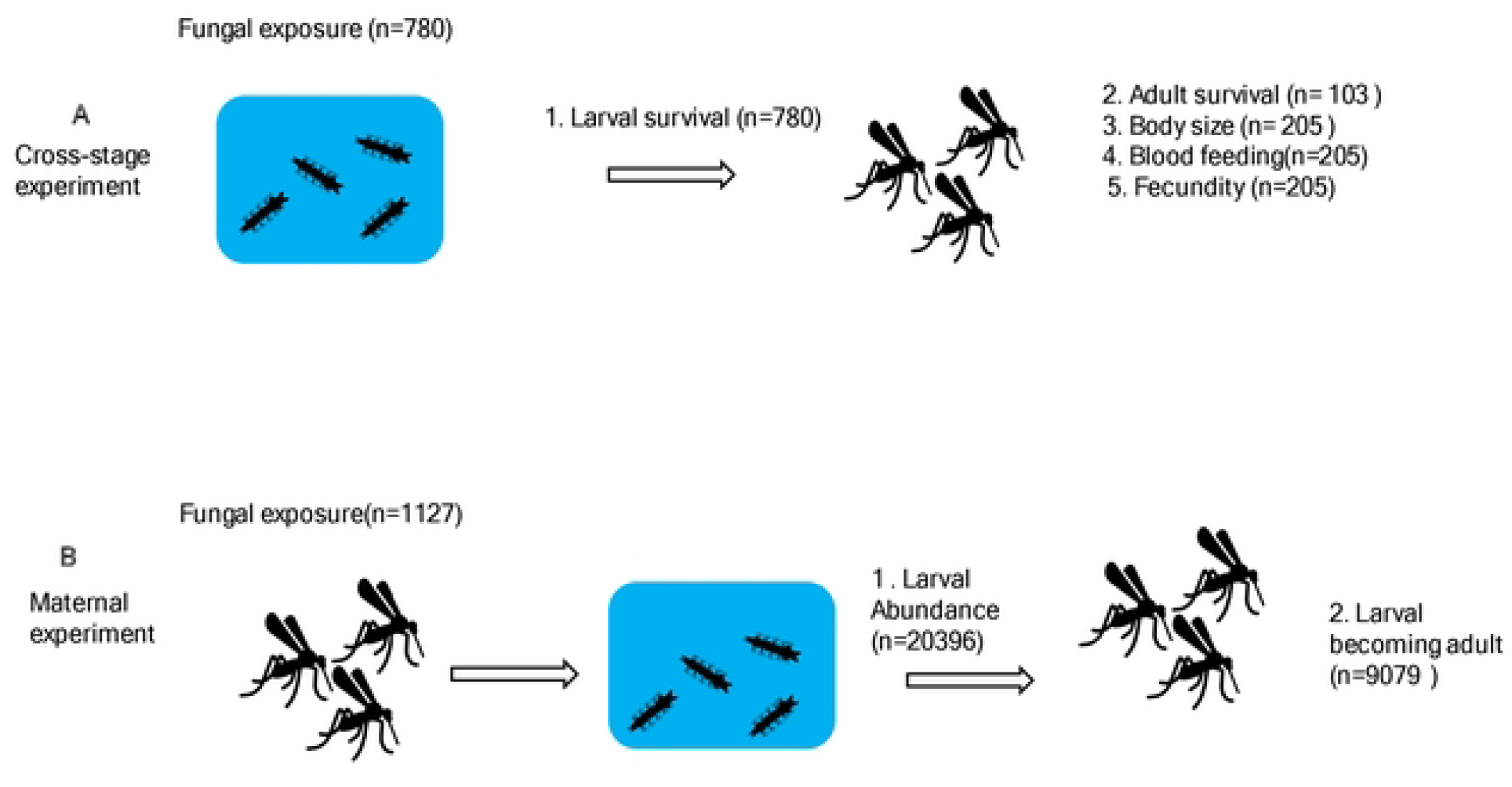
Experimental graphical design. In the cross-stage experiment (A) larvae were exposed to fungi or control, then larval survival and emerging adult traits (survival, body size, blood feeding and fecundity) measured. In the maternal experiment, (B) female adult mosquitoes were exposed to fungi or control and the number of larvae and emerging adults measured.

### Fungal suspension preparation

Using a sterile spatula, small fragments of fungal mycelium and conidia were carefully collected by gently scraping the surface of the fungal culture in a Petri dish under aseptic conditions to prevent contamination[23,25]. The harvested material was then suspended in a sterile aqueous solution containing 0.05% Tween 80, a non-ionic surfactant commonly used to improve conidial dispersion and prevent clumping. This suspension was vortexed for several minutes to ensure a homogeneous distribution of conidia.

To determine the conidial concentration, an aliquot of the suspension was subjected to quantification using a Neubauer hemocytometer under a phase-contrast microscope. The desired concentration of 10^7^ conidia/mL was achieved by serial dilution or concentration adjustment, ensuring a standardized inoculum density for bioassays.

Once prepared, the conidial suspension was introduced into larval breeding trays containing third and fourth instar mosquito larvae. The trays were maintained under controlled environmental conditions, including temperature (27 ± 2°C), relative humidity (75 ± 5%), and a 12:12 h light-dark photoperiod, to mimic natural breeding habitats. The exposure period was standardized to allow sufficient fungal attachment and germination on larval cuticles, facilitating infection. This experimental setup enabled the assessment of fungal pathogenicity and virulence against mosquito larvae under laboratory conditions.

### Mosquito rearing

The mosquito strain of *Anopheles coluzzii* used in the experiment was the 11^th^ generation of a line that originated from the Vallée du Kou and was established in the laboratory at Institut de Recherche en Sciences de la Santé (IRSS), Bobo Dioulasso, Burkina Faso. The colony was maintained at 27 ± 2°C, relative humidity of 70 ± 5% and photoperiod of 12L:12D.This colony is known to have almost fixed 1014F *Kdr* allele [26]. The larvae were kept in plastic trays filled with tap water and fed at all stages with Tetra-min^®^. All emerged mosquitoes had access to 6% glucose and female mosquitoes were fed on rabbit blood. Eggs were laid on wet filter paper in the cages and transferred to the larval trays.

### Larval survival after fungal infection

A total of ten plastic trays were used in the experiment, with each tray containing 30 L4 larvae. This setup was replicated three times, resulting in a total of 780 larvae. The larvae were exposed to the fungal solution and remained in the treated environment until they reached the pupal stage. Each tray contained a total volume of 50 mL, composed of 49 mL of tap water and 1 mL of a fungal suspension at an initial concentration of 10^7^conidia/mL. The addition of the fungal solution to the water led to a final concentration of 2×10^5^ conidia/mL in each tray. Then, we monitored and recorded larval survival rates throughout their development, from the larval stage to adulthood.

### Survival of emerging adults

The adult mosquitoes that successfully emerged from the treated larval suspension were carefully collected and transferred to designated rearing cages(~ 30 mosquitoes per cage) under controlled conditions. Inside the cages, they were provided with a continuous supply of a 5% glucose solution to ensure proper feeding. Mortality was monitored daily, with dead mosquitoes being systematically removed from the cages for up to five days post-emergence. This allowed for the assessment of delayed mortality effects potentially caused by fungal exposure during the larval stage.

### Mosquitoes wing size measuring after fungal infection

The left wings of the adult mosquitoes were carefully dissected using fine-tipped forceps and a sterile needle to ensure precision and minimize structural damage. Each excised wing was then mounted onto a microscope slide with a coverslip for detailed morphometric analysis, following the methodology described by[27].

High-resolution digital images of the mounted wings were captured using a Nikon SMZ1500 stereomicroscope (Nikon, Japan) equipped with an integrated camera. The imaging process was conducted at a magnification of 11.25X to ensure accurate measurement of wing dimensions. Wing length was determined by measuring the linear distance from the distal wing tip to the alular notch, a standard landmark for wing morphometry in mosquitoes.

A total of 225 mosquitoes were analyzed for this experiment, with the study conducted in three independent replicates to ensure statistical robustness and reproducibility of the findings.

### Fungus infection effect on female mosquito fecundity and fertility

Four experimental cages, each containing approximately 100 mosquitoes (50 males and 50 females), were used to assess the impact of fungal exposure on mosquito reproductive success. The mosquitoes, aged 3–5 days post-emergence, originated from either the fungal-treated larval suspension or the untreated control group. To ensure sufficient blood intake for egg development, all mosquitoes were offered a rabbit blood meal twice, thereby maximizing the likelihood of successful feeding.

Following blood feeding, engorged females were individually transferred to oviposition cups to facilitate precise monitoring of egg-laying behavior. The total number of eggs laid per female was recorded to evaluate fecundity. A total of 205 females were analyzed for this experiment.

To assess egg viability, the collected eggs were submerged in 50 mL of tap water, and their hatching success was systematically evaluated. This step allowed for the determination of potential carryover effects of fungal exposure on mosquito reproductive output and offspring development.

### Effect of fungal infection of adult mosquitoes on their offspring (G1)

For each treatment, approximately 100 blood-fed female mosquitoes were exposed to fungal suspensions (S10 and S26) in 10 independent replicates. Prior to fungal application, mosquitoes were temporarily immobilized by chilling in a freezer at –4°C for 15 seconds, ensuring minimal stress while facilitating uniform exposure to fungal spores. Immobilized mosquitoes were then transferred onto a Petri dish lined with sterile filter paper to maintain aseptic conditions and prevent contamination during the spraying process.

Mosquitoes were sprayed with a 1 mL fungal suspension containing *M. pingshaense* conidia at a standardized concentration of 1×10^7^ spores/mL, formulated in 0.01% (v/v) aqueous Tween 80. This formulation was applied using an Ami pulvérisateur (Zhejiang, China), a precision spray device designed to ensure consistent and homogeneous deposition of conidia on mosquito cuticles. Control mosquitoes were sprayed with 1 mL of 0.01% (v/v) aqueous Tween 80 alone, serving as a negative control to account for any effects of the spraying procedure.After treatment, mosquitoes were transferred to holding cups (7 cm diameter × 9 cm height) and maintained under controlled environmental conditions: temperature of 25°C, relative humidity of 70 ± 10%, and a 12:12-hour light-dark cycle. To sustain physiological activity, mosquitoes were provided a 5% glucose solution soaked in a cotton ball. Egg-laying behavior was monitored, and the number of fourth-stage larvae (L4) and the number of larvae successfully emerging as adults were recorded.

## Statistical analysis

Larval survival(**figure 2**) rate was analyzed using a binomial Generalized Linear Model. Treatment (3 levels: Control, S10 and S26), time (4 levels: J1, J2, J3 and J4) and their interaction were included as fixed effects. A Cox proportional hazard models from the R package “survival”, was developed to determine the impact of fungal solution on the survival of adults (**Figure 3 A**). For this, adult survival was the response variable with treatment was included as fixed effect and the random effect of ‘replicate’ was incorporated as a frailty function[28,29]. To understand the impact of the treatment on different entomological parameters of the adult mosquitoes, separate generalized linear mixed models (GLMMs) with negative binomial family distribution were developed with the following response variables: i)feeding proportion (Figure 3B); ii) number of eggs (**Figure 3C**); iii) Wing size (**Figure 3 D&E**), vi) hatched larvae (**Figure 4A**); v) generation G1(first offsprings from infected mothers) (**Figure 4B**) using the R package ‘glmmTMB’. Other parameters such as larval abundance was modelled using a negative binomial family distributed to account for the full dispersal in the data, while wing size was modelled following a Gaussian distributed GLMM with R package “lme4”. All these models were fitted with treatment as fixed effect and replicate as random effect. For all models described above we performed model selection using stepwise removal of terms, followed by likelihood ratio tests. The best model retained only significant terms that improved model goodness. Model performance diagnostics (i.e., residuals and dispersion) were evaluated for all models using the R package ‘DHARma’. All statistical analysis were performed using R version 4.1.2.

**Figure 2:**
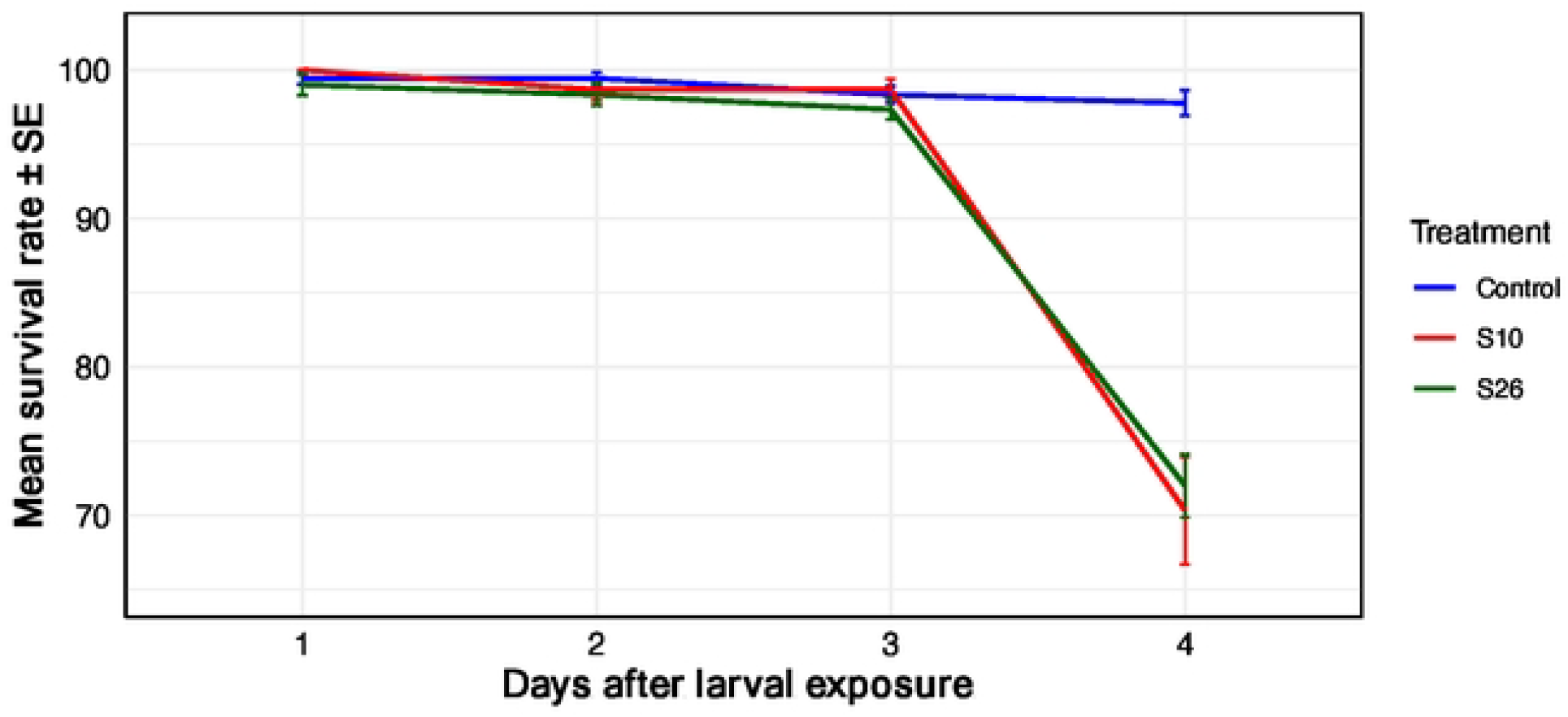
Cross-stage experiment, fungal effect on larval survival. *Temporal changes in mosquito larval survival under different treatments*. Mean survival rates (± standard error) of 4^th^-instar larvae exposed to fungal isolates S10 (in red) and S26 (in green) compared to the untreated control (in blue) over a four-day period. Each point represents the average survival rate for a treatment group at a given time point. Lines connect points to show temporal trends, and error bars indicate standard error of the mean.

**Figure 3:**
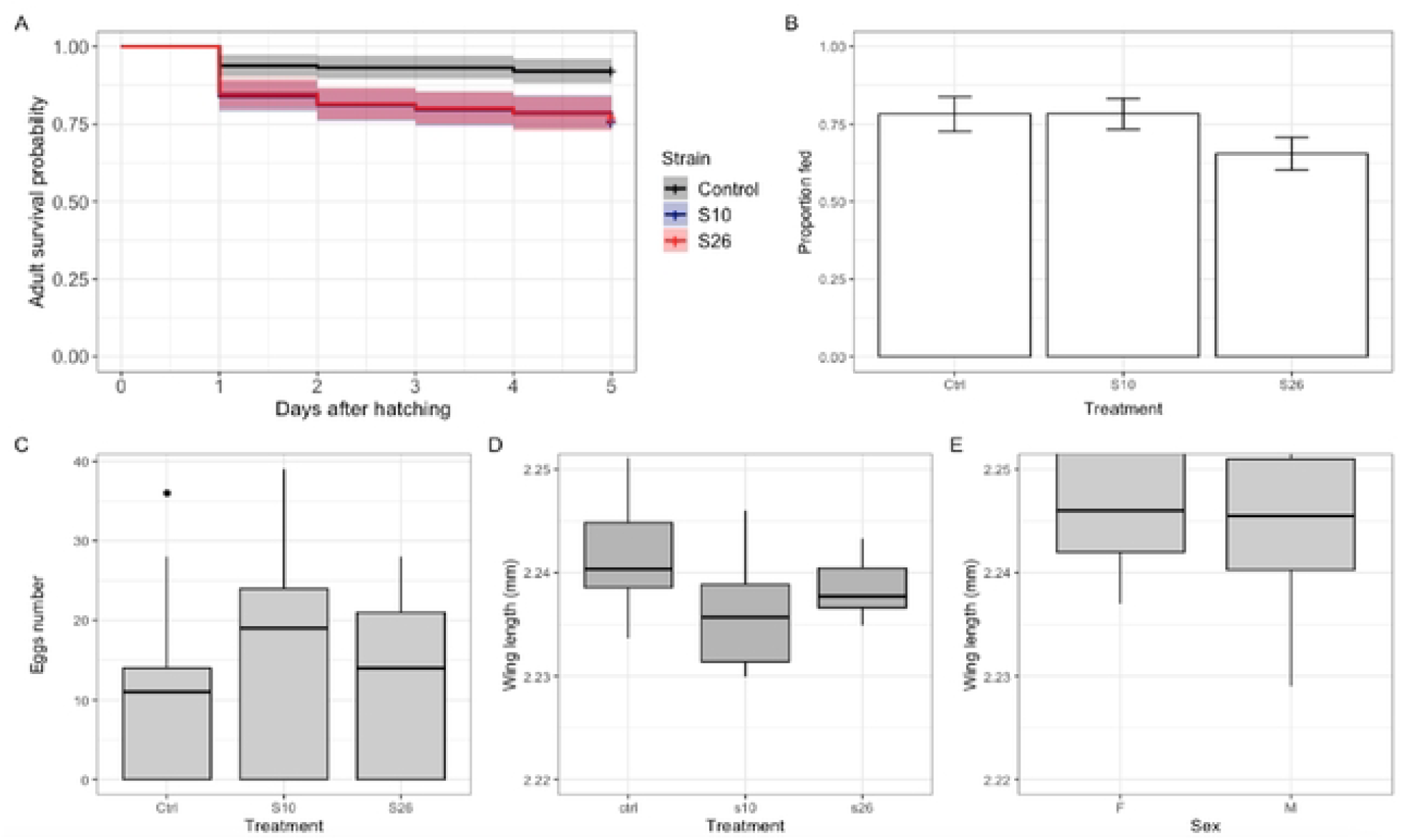
Cross-stage experiment, fungal effect on adult life-history traits. **(A)** Kaplan Meier survival curves of mosquito larvae exposed to fungal solution (S10 blue, S26 in red) and a control group (black). Solid lines represent predicted mean survival and shaded areas represent 95% confidence intervals. The plus sign indicates censored data. Shaded areas represent 95% confidence interval. **(B)** The proportion of female adults surviving fungal exposure that took a blood meal and **(C)** the number of eggs that they laid. **(D)** Impact of fungal infection on mosquito size, (**E**) Impact of fungal infection on mosquito size based on sex.

**Figure 4:**
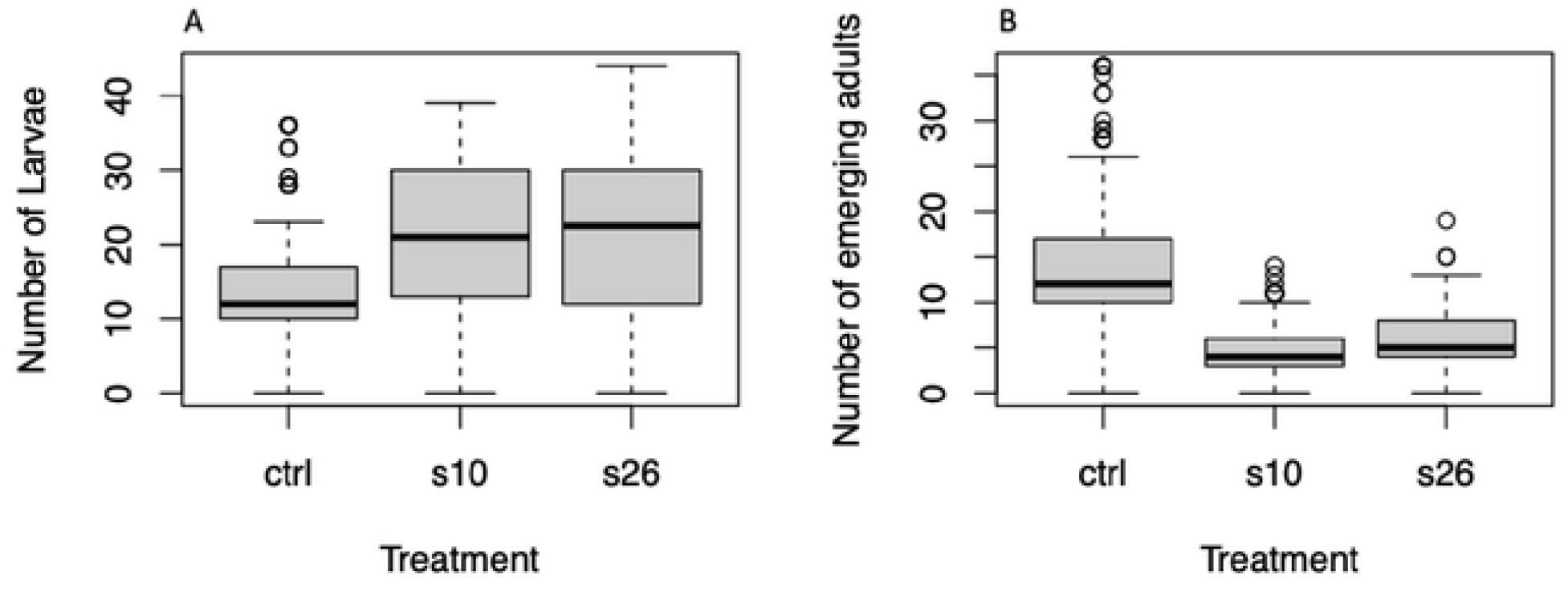
Maternal Experiment, fungal effect on progeny. The different effects of the fungus on the life traits of mosquitoes. larvae hatched when fungal infection occurs during the adult stage **(A);** The impact of fungal infection on larvae abundance **(B)**.

## Results

### Exposure of larvae to fungi increases mortality compared to controls

There was a significant treatment × time interaction effect on larval survival rate (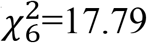, *p*=0.006); At J1, J2 and J3(days post exposure), larval survival (L4 stage) rate was close to 100% regardless of treatment; however, at J4, larval survival rate in the control treatment batch was still close to 100% while it was down to approx. 70% in both the S10 and S26 fungal treatment batches (**Figure2**).

### Fungal exposure at larval stage reduces survival of emerging adults and increases their fecundity

We monitored the survival of the 103 adults that emerged after being exposed to fungi or control at the larval stage for 5 days after emergence. We found that fungal exposure decreased adult mosquitoes’ survival (*X*^*2*^ *= 26*.*361, df =2, p< 0*.*001*) (**Figure 3A)**.

Female adults were blood fed to determine the impact of infection on feeding behavior and fecundity. We analysed a total of 205 female mosquitoes and found that fungal treatment during the larval stage did not influence the proportion of blood fed females (*X*^*2*^*= 96*.*434, df = 2*, p= 0.131) (**Figure 3B**).

After removing non-blood fed mosquitoes, we monitored egg-laying in 150 fed mosquitoes individually and found that fungal treatment during the larval stage increased the number of eggs laid compared to controls by 1.27 times (*X*^*2*^*= 96*.*434, df = 2, p<0*.*001*) (**Figure 3C**).

### Larval fungal exposure does not alter the body size of emerged adults compared to control

The increased number of eggs laid by adults emerging from larval fungal infection suggests that infection might have resulted in larger adults, which is associated with increased fecundity [30]. To test this hypothesis, we measured mosquito wing lengths as a proxy of their body size between control and fungal infection treatments. Surprisingly, we found that fungal treatment at the larval stage had no effect on mosquito size (*X*^*2*^*=1*.*0292, df =2, p=0*.*5977)* **(Figure 3D)**, suggesting that the fecundity effect is mediated by the direct effect of the fungi, not on selection of larger individuals. Overall, females were larger than males (*X*^*2*^*= 3*.*8963, df =1, p= 0*.*04839*) **(Figure 3E)**.

### Maternal fungal exposure increases the abundance of larval progeny but decreases the number of emerging adults compared to control

As we found cross-stages effects of fungal exposure, where treatment of larvae decreased adult survival and increased fecundity, we tested if infection at the maternal stage could also affect the progeny by infecting adult females and measuring larval and adult abundance of the progeny. Out of the total 266 female mosquitoes treated, we found that maternal fungal treatment significantly increased the number of larvae (L4) compared to controls. Indeed, the number of mosquitoes larvae treated with both S26 and S10 was 1.44 and 1.47 times higher, respectively, than the untreated control *(X*^*2*^*= 654*.*57,df =2, p<0*.*001)* (**Figure 4A)**. However, the number of larvae that became adults was two times lower in the groups treated with the fungi than in the untreated ones (X^2^= 642.49, df =2, p<0.001) **(Figure 4B**).

## Discussion

Efficient management of mosquito breeding sites would be a powerful complementary tool for the control of malaria. This could involve the use of natural enemies, including entomopathogens such as *Metharizium*. This study confirmed previous findings that *Metharizium* can be effective at targeting malaria mosquito larval stages[11,31] and added novel insights into the cross-stage and generation effects. We found that although larvae that are exposed to fungi have a lower survival rate, those that survive as emerged adults generally had increased mortality but their fecundity was increased in females. Together our findings show there are some trade-offs of the impact of *Metarhizium* exposure on different life-history traits.

We found that infecting adults mosquitoes increase the number of larvae produced in the treated population compared to the control population. This suggests that the infected individuals are under enormous pressure to survive and they are forced to invest in early reproduction. This finding is in line with studies [32–34] that reported the entomopathogenic fungi *Beauveria bassiana* and *Metarhizium anisopliae* significantly reduce fecundity and egg viability in various hosts, thereby compromising their offspring. Their impact goes beyond direct mortality by exerting reproductive pressure, enhancing their potential as biological control agents. However, we noted that a very small proportion of these larvae reach the adult stage in the next generation. Since the larval density was standardized across treatments, we believe the increased mortality cannot be explain by density pressure. Instead, a plausible explanation could be that the eggs were immature, so those that hatched did not possess the necessary biological components for proper development. Also, regarding the infection at larval stage the results show how certain using of entomopathogenic fungus can improve the ability of spores to spread on a water surface and infect their hosts. We found high mortality of the larvae in both treatments at the pupal stage. This suggests that the larvae ingested a lethal dose of the fungus as food during their larval stage similar results have been reported by [35–39]. That studies reported that high pupal-stage mortality suggests larvae ingested lethal fungal doses during development.

Adults from larvae exposed to fungi solution had a relatively low survival rate compared to control group. We hypothesize that the spore had already passed through the larval cuticle before moulting into the adult stage[20,40,41], so these spores expressed their toxins once in the adult mosquito’s haemolymph or that, even in the larval stage, the fungus had already passed through the mosquito’s haemolymph. Indeed, as *An. coluzzii* larvae have different rates of filtration and ingestion on the surface and the spore can infect by ingestion or contact [16]; However, for those larvae that survived the infection, it is possible that the spore was still on the surface of the host and that during molting the spore was shed or that there was no contact between the larva and the spore[40,41]. Molting has been reported to be an important factor in the resistance of arthropods to fungal infection, particularly in arthropods with short ecdysis intervals [29].

In a context where the multiplicity of mosquito breeding sites is a real bottleneck for malaria control programs. The use of fungi in larval breeding sites could help to reduce mosquito populations at both the larval and adult stages[42].

Observation of the size and sex of treated and untreated individuals showed no significant difference. This is consistent with previous studies showing similar wing size measurements between untreated and treated individuals with the fungi, but they show males mosquitoes Were bigger than female’s [27][43]. This suggests that infection at the larval stage does not influence the size of individuals according to their treatment status; however, females appear larger than males emerging from the fungal solution [16,19,38]. Based on the data collected regarding larval exposure to entomopathogenic fungi, the results can be interpreted as follows: infected larvae tend to occur at lower densities within their habitat, which may promote enhanced individual growth and, consequently, lead to higher egg production at the adult stage. However, this hypothesis is not supported by body size measurements, which did not reveal significant differences between infected and control individuals. This suggests that the observed increase in fecundity is not mediated by body size variation but rather by a physiological response triggered by the infection. It is plausible that the physiological stress induced by fungal exposure prompts females to invest more heavily in early reproduction, possibly as a compensatory mechanism in anticipation of reduced lifespan. This interpretation is consistent with observations made in infected adults, where a higher number of eggs laid was also recorded, reinforcing the hypothesis of increased reproductive effort in infected or stressed females.These findings confirm that entomopathogenic fungi and their effective effects are possible candidates to swap synthetic insecticides for controlling larvae, pupae and adult mosquitoes [44,45].

## Conclusion

*Metarhizium* seems to be a promising biocontrol agent for many insects including mosquitoes. Although feeding, oviposition and mosquito size do not seem to be influenced by the fungi treatment at the concentration tested, *An. coluzzi* lifespan seems to be greatly reduced when they are infected with conidia and the fungal spores seem to also impact the survival of larvae and adult emergence and survival, which will likely have important consequences for vectorial capacity. These findings highlight the complex trade-offs induced by fungal infection and support the integration of entomopathogens into vector control strategies as a complementary or alternative tool to synthetic insecticides.

## Acknowledgements

We express our sincere gratitude to all the study participants for their time and contribution to this study. We are grateful to Vallée du Kou community and IRSS lab technicians du for their help in conducting lab activities.

## Author contributions

IS, FB, AB, FD, MV,AD, AM, LL, AT and EB conceived of the study. IS, EB conducted the experiments. FB, MV, IS and EB analysed the data. All authors drafted the manuscript. All authors read and approved the final manuscript.

## Funding

This work was supported by the National Institute for Health Research (NIHR) (using the UK’s Official Development Assistance (ODA) Funding) and Wellcome Trust grant ref218771/Z/19/Z under the NIHR-Wellcome Partnership for Global Health Research. The views expressed are those of the authors and not necessarily those of Wellcome Trust, the NIHR or the Department of Health and Social Care’. Preliminary field activities were supported by the Open Philanthropy grant. FB is supported by the Academy Medical Sciences Springboard Award (ref:SBF007\100094). MV is supported by the European Research Council under the European Union’s Horizon 2020 Research and Innovation Programme (grant agreement no. 852957)

## Ethical approval

Not applicable

## Availability of data and materials

All data for this study will be available upon request.

## Abbreviations

B.: *Beauveria*
*An.*: *Anopheles*
*Crtl*: *control group*
*S10*: *Metarhizium* strain 10 in our stump library
*S26*: *Metarhizium* strain 26 in our stump library

## Competing interests

The authors declare no competing interests.

## Notes

### Competing Interest Statement

The authors have declared no competing interest.

